# Quantifying how constraints limit the diversity of viable routes to adaptation

**DOI:** 10.1101/279661

**Authors:** Sam Yeaman, Aleeza C. Gerstein, Kathryn A. Hodgins, Michael C. Whitlock

## Abstract

Convergent adaptation can occur at the genome scale when independently evolving lineages use the same genes to respond to similar selection pressures. These patterns provide insights into the factors that facilitate or constrain the diversity of genetic responses that contribute to adaptive evolution. A first step in studying such factors is to quantify the observed amount of repeatability relative to expectations under a null hypothesis. Here, we formulate a novel metric to quantify the constraints driving the observed amount of repeated adaptation in pairwise contrasts based on the hypergeometric distribution, and then generalize this for simultaneous analysis of multiple lineages. This metric is explicitly based on the probability of observing a given amount of repeatability by chance under an arbitrary null hypothesis, and is readily compared among different species and types of trait. We also formulate a metric to quantify the effective proportion of genes in the genome that have the potential to contribute to adaptation. As an example of how these metrics can be used to draw inferences, we assess the amount of repeatability observed in existing datasets on adaptation to antibiotics in yeast and climate in conifers. This approach provides a method to test a wide range of hypotheses about how different kinds of factors can facilitate or constrain the diversity of genetic responses observed during adaptive evolution.

## Introduction

What factors limit the diversity of viable genetic routes to adaptation? If different species encounter the same selection pressure, will adaptive responses occur through mutations in homologous nucleotides, regulatory regions, protein domains, genes, or gene pathways? Empirical studies have identified different amounts of convergent adaptation across a range of species, traits, timescales, and levels of genetic hierarchy [1–3]. But why do similar forms evolve at the genomic level, and what does the level of convergence tell us about underlying constraints? Does evolution use the same genes repeatedly because there are only a limited number of ways that genetic and developmental pathways can generate a given phenotype or because only a limited proportion of generated phenotypes are selectively optimal? These two explanations represent fundamentally different kinds of constraints affecting the diversity of genetic responses generated by evolution, so studying their relative importance is central to understanding adaptation. Our broad aim here is to develop methods that represent how empirical observations deviate from null hypotheses under different models of the mapping of genotype to phenotype to fitness. While discriminating between these alternative models may prove difficult in practice, this provides a first step towards quantifying constraints in a way that is readily compared across study systems, with the eventual goal being a comprehensive understanding of the relative importance of different factors shaping the diversity of routes to adaptation.

To rephrase the above questions in quantitative terms based on the flexibility of the mapping from genotype to phenotype to fitness: does repeatability occur because of low redundancy in the mapping of genotype to phenotype (only a few ways to make the same phenotype; Figure 1A), or because of low redundancy in the mapping of genotype to fitness? (only a subset of the genotypes yielding the same phenotype are optimal; Figure 1B). Redundancy in the mapping of genotype to phenotype (hereafter, GP- redundancy; [4]) is determined by two factors: 1) the difference between the number of genes that need to mutate to yield a given phenotype and the number of genes that could potentially mutate to give rise to variation in the trait, and 2) the extent to which different genes have interchangeable vs. uniquely important effects on the phenotype. High GP- redundancy means that many different combinations of alleles can have the same phenotype, so if all else is equal, then independent bouts of adaptation are likely to occur via different sets of mutations and repeatability will be low ([4,5]). The standard quantitative genetic model implicitly assumes complete GP-redundancy with fully interchangeable allelic effects, while the recently proposed omnigenic model assumes high but incomplete redundancy, with “core” vs. “peripheral” genes having different potential to affect variation [6].

**Figure 1.**
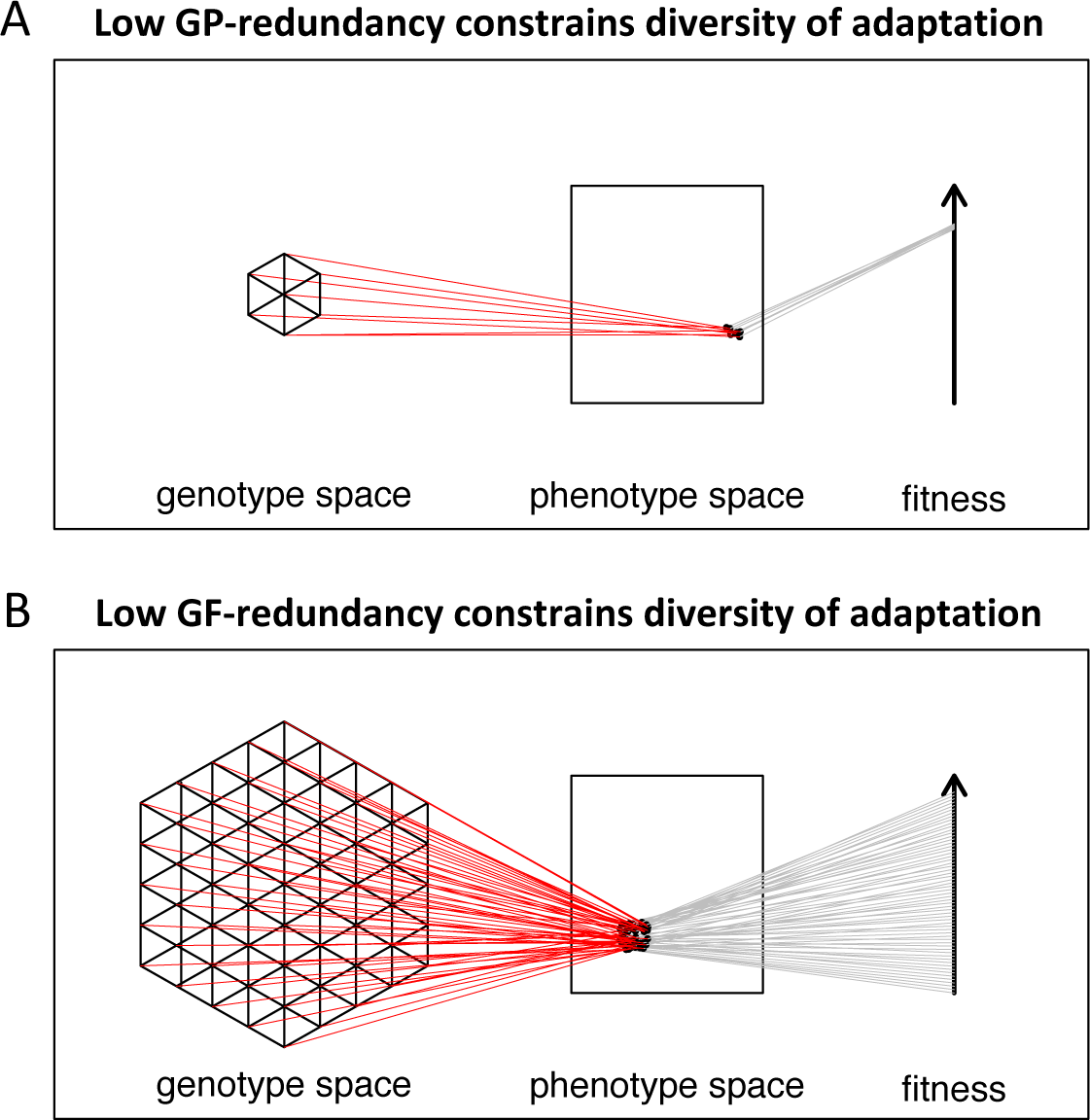
Scenarios with different combinations of GP- and GF-redundancy that result in high repeatability of adaptation (adapted from [14]).

Redundancy in the mapping of genotype to fitness (hereafter, GF-redundancy) is determined by both the direct effects of genotype on phenotype and phenotype on fitness and a number of possible indirect effects. These indirect effects can arise due to epistatic interactions or pleiotropic effects on other selected traits. Alternatively, GF-redundancy can also be affected by aspects of the genetic architecture such as the number of alleles and their linkage relationships and effect sizes, depending upon the interaction between migration, selection, and drift. For example, if a given phenotype is coded by many small unlinked alleles, this architecture would be less fit than a similar phenotype coded by a few large or tightly linked alleles, in the context of migration-selection balance [7] or negative frequency dependence [8,9]. Similarly, the increased drift that occurs in small populations may prevent alleles of small effect from responding to natural selection [10,11], resulting in such genotypes being effectively neutral and therefore lower in realized fitness than those made up of large-effect alleles. Polygenic models of directional selection (*e.g*. [12]) assume no GP- and GF-redundancy, while traditional quantitative genetic models of Gaussian stabilizing selection assume high GP- and GF- redundancy (*e.g*. [13]). Rephrasing the main question in terms of redundancy: when high repeatability is observed, is the low diversity of genetic routes to adaptation being constrained by low GP-redundancy (few ways to make a phenotype) or by high GP- redundancy but low GF-redundancy (many ways but few are good)?

In addition to these types of redundancy, differences in mutation rate among genes or types of variants (*e.g*. SNPs, indels, microsatellites) can also greatly affect the repeatability of adaptation [2,15–17]. Genes that can mutate to a beneficial phenotype through loss-of-function mutations are often implicated in repeated evolution (e.g. [18,19]), likely because there are more ways to break a gene than to beneficially refine its protein function. Also, in cases where there is insufficient time since the most recent common ancestor for complete lineage sorting, shared standing variation can greatly increase repeatability, due to higher fixation probability of variants at intermediate frequency [17,20]. Finally, if some genes tend to maintain more standing variation than other genes, then repeatability will be higher for genes with higher standing variation, even if the actual causal alleles are different in each lineage. If a given gene has the capacity to mutate more rapidly or maintain more standing variation, these factors could be seen as fundamental drivers of evolvability that contribute to realized to GP- and GF- redundancy. However, because shared standing variation is more dependent on historical contingency, it is important to control for its contribution when drawing inferences about the importance of redundancy (see Discussion).

It is important to note that the constraint that we discuss here is only referring to factors that affect the diversity of genes used in independent bouts of adaptation, rather than factors that limit an adaptive phenotypic response in general. Observing the same gene contribute to adaptation in numerous lineages (*e.g. Mc1r*; [21]) can rightly be interpreted as evidence that some feature of the interaction between the developmental-genetic program and ecology facilitates the rapid emergence of adaptation, rather than constraining it. However, the same example can also be interpreted as being severely constrained in terms of the diversity of forms, since there are so few viable alternative genetic solutions that actually evolve [19]. Thus, evolutionary constraints can be considered along two related but distinct axes: factors that affect the potential for any adaptive response [22,23] vs. factors that affect the diversity of forms (*i.e*., genetic routes to adaptation). A scenario that is highly constrained in terms of the diversity of forms may be least constrained in terms of the potential for a rapid adaptive response to a change in environment, and low redundancy in the mapping of genotype-phenotype-fitness may itself be a product of adaptation over deep time. For the remainder of this manuscript, we focus on constraints affecting the diversity of forms, which we refer to as “diversity constraints” for simplicity.

There has been considerable discussion in the literature about the effects of these different factors on convergence [3,16,17,20,24–28], and various metrics have been used to quantify repeatability in empirical contexts (*e.g*. Jaccard index, Proportional Similarity; [2,1]). However, there are several limitations in using these metrics to study diversity constraints, as they do not incorporate information about genes that could contribute to adaptation but don’t and are not explicitly tied to the probability of repeatability occurring under a null model. These existing metrics provide a useful description of how often the same gene is used in adaptation, but as we will show below, they are not well-suited for testing of hypotheses to discriminate between these different kinds of constraint.

Here, we develop statistical approaches for quantifying the diversity constraints that drive repeatability in genomic data from studies of local adaptation and experimental evolution. To study these constraints, we formulate an explicit probability-based representation of the deviation of observed repeatability from expectations under different null hypotheses. This approach can be used after standard tests have been applied to identify the putative genes driving adaptation, and uses as input either binary categorization of genes as “adapted” or “non-adapted” or any continuous metric representing the relative amount of evidence for a given gene being involved in adaptation (*e.g. F*_ST_, *p*-values, Bayes factors). We begin by formulating an analytical model for a contrast of two lineages with binary data, and then generalize this model for contrasts of multiple lineages using either binary or continuous data. We also propose a novel metric estimating the proportion of genes in the genome that can potentially give rise to adaptation. In all cases, these models can be used to successively test null hypotheses that incorporate different amounts of information about the constraints that could shape repeatability.

The simplest null hypothesis is that there are no constraints and all genes have equal probability of contributing to adaptation. If more repeatability is observed than expected under this null model, then two inferences can be made: natural selection is driving patterns of convergence (and that observed signatures are not false positives), and some diversity constraints are operating to increase the repeatability of adaptation. We then consider how other null hypotheses can be formulated to represent the various kinds of constraints discussed above. We focus mainly on the effect of low GP-redundancy, where the number of genes that could potentially contribute to adaptation is much smaller than the total number of genes in the genome, but also discuss how constraints arising from GF-redundancy, standing variation, or mutation rate could be modeled. Because this method quantifies repeatability in terms of probability-scaled deviations from expectations, it can be applied across any trait or species of interest, allowing contrasts to be made on the same scale of measurement.

## Methods

### Quantifying diversity constraints in pairwise contrasts

Suppose there are two lineages, *x* and *y*, that have recently undergone adaptation to a given selection pressure, resulting in convergent evolution of the same phenotype within each lineage. This adaptation could be global, with new mutations fixed within lineages (*e.g*., in experimental evolution studies with multiple replicate populations), or local, with mutations contributing to divergence among populations within each lineage (*e.g.*, in observational studies of natural adaptation to environmental gradients). In either case, we assume that adaptation can be reduced to a binary categorization of genes as “adapted” or “non-adapted”. We use the following notation to represent different properties of the genomic basis of trait variation: the number of loci in the genome of each species is *n*_*x*_, and *n*_*y*_, with the number of orthologous loci shared by both species being *n*_*s*_; the adaptive trait is controlled by *g*_*x*_ and *g*_*y*_ loci in each species, with *g*_*s*_ shared loci (*i.e*. the loci in which mutations will give rise to phenotypic variation in the trait, hereafter the “mutational target”); of the *g* loci that give rise to variation, only a subset have the potential to contribute to adaptation due to the combined effect of all constraints, represented by *ga*_*x*_ and *ga*_*y*_, with *ga*_*s*_ shared loci (the “effective adaptive target”); in a given bout of adaptation, the number of loci that contribute to adaptation in each lineage is *a*_*x*_ and *a*_*y*_, with *a*_*s*_ orthologous loci contributing in both lineages. For simplicity, we assume that there is complete overlap in the genomes (*n*_*s*_ = *n*_*x*_ = *n*_*y*_), mutational targets (*g*_*s*_ = *g*_*x*_ = *g*_*y*_), and loci potentially contributing to adaptation (*ga*_*s*_ = *ga*_*x*_ = *ga*_*y*_) in both species (see supplementary materials and Figure S1 for set notation). These assumptions are most appropriate for lineages that are relatively recently diverged, where most orthologous genes are retained at the same copy number and the developmental-genetic program is relatively conserved, so that the same genes potentially give rise to variation in both lineages. Lineages separated by greater amounts of time would be expected to have reduced *n*_*s*_ due to gene deletion, duplication, and pseudogenization in either lineage, and reduced *g*_*s*_ and *ga*_*s*_ due to evolution and divergence of the developmental-genetic program, through sub- and neo-functionalization, and divergence in regulatory networks.

Under the assumption that all *ga*_*s*_ genes have equal probability of contributing to adaptation, the amount over overlap in the complement of genes that are adapted in both lineages (*a*_*s*_) is described by a hypergeometric distribution where the expected amount of overlap is *ā*_*s*_ = *a*_*x*_*a*_*y*_*/ga*_*s*_ (*e.g*. [29]). In practice, we typically have little prior knowledge about which genes have the potential to contribute to either adaptation (*ga*_*s*_) or standing variation in the trait (*g*_*s*_), but we can draw inferences about how these parameters constrain the diversity of adaptive responses by testing hypotheses and comparing the observed amount of overlap (*a*_*s*_) to the amount expected under a given null hypothesis (*ā* _*s*_). To test different hypotheses about how diversity constraints give rise to repeated adaptation, we represent the total number of genes included in the test set as *g*_0_. The simplest null hypothesis is that there are no diversity constraints and all genes potentially give rise to variation and contribute to adaptation (*g*_0_ = *ga*_*s*_ = *g*_*s*_ = *n*_*s*_), so by rejecting this null, we can infer that *ga*_*s*_ < *n*_*s*_. In model systems where something is known about which genes potentially contribute to variation for the trait (based on mutation accumulation or GWAS), then a more refined null hypothesis can be tested, where *g*_0_ = *g*_*s*_. By rejecting this null, we can infer that *ga*_*s*_ < *g*_*s*_, which could occur due to low GF-redundancy or differences among genes in mutation rate or standing variation. We can also reverse the direction of inquiry and estimate *ga*_*s*_ directly from the data by calculating 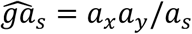, such that a metric representing the effective proportion of the genome that can potentially contribute to adaptation can be calculated as 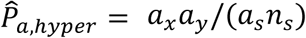

For any value of *g*_0_, an effect size representing the excess in overlap due to convergence relative to the null hypothesis can be expressed by standardizing the observed overlap by subtracting the mean (*ā*_*s*_ = *a*_*x*_*a*_*y*_*/g*_*0*_) and dividing by the standard deviation of the hypergeometric distribution:

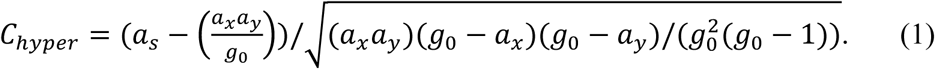

This metric provides a quantitative representation of how much more overlap occurs than expected under the null hypothesis, scaled according to how much a given bout of evolution would deviate from this expectation if the null hypothesis were true. Similarly, the exact probability of observing *a*_*s*_ or more shared loci contributing to adaptation can also be calculated using the hypergeometric probability (see Supplementary Information for sample R-script), which provides a *p*-value.

### Quantifying diversity constraints in multiple lineages

While pairwise contrasts are most straightforward statistically, they have considerably lower power than comparisons among multiple lineages. If one gene (such as *Mc1r*) tends to drive adaptation repeatedly in a large number of lineages, this may go undetected in an approach using multiple pairwise comparisons, but would be readily detected in a simultaneous comparison of multiple lineages. Unfortunately, while the hypergeometric distribution provides an exact analytical prediction for the amount of overlap in a pairwise comparison, which can be used to calculate a *p*-value and the probability-based effect size (*C*_*hyper*_), it cannot be easily generalized to simultaneously analyze multiple lineages. While it is possible to conduct pairwise analysis and average the results across multiple comparisons, *p*-values from this approach might fail to detect cases where a single gene contributes repeatedly to adaptation in more than two lineages, as information does not transfer among the pairwise comparisons. We now develop an alternate, approximate approach to assess repeatability in multiple lineages by calculating Pearson’s *χ*^*2*^ goodness of fit statistic and comparing this to a null distribution of *χ*^*2*^ statistics simulated under the null hypothesis to calculate a *p*-value and an effect size. The *p*-value obtained by this approach represents the probability of observing a test statistic as extreme or more extreme under the null hypothesis, considering all lineages simultaneously. The effect size is instead calculated as an average across all pairwise comparisons among the *k* replicate lineages, so that it represents the increase in repeatability relative to expectations under the null for a given bout of adaptation in a single lineage (and does not therefore depend upon sampling effort in terms of the number of lineages).

Consider the case where *g*_0_ genes can potentially contribute to adaptation in the given trait and each lineage has some complement of genes that have mutated to drive adaptation, with *α*_*i,j*_ representing the binary score for gene *i* in lineage *j* (1 = adapted, 0 = non-adapted). The summation for gene *i* across all lineages provides the observed counts (*o*_*i*_ = Σ_*j*_ *α*_*i,j*_) while the expected counts (*e*_i_) can be set based on the null hypothesis being tested. Under null hypotheses where all genes in *g*_0_ have equal probability of contributing to adaptation, the expected counts are equal to the mean of the observed counts (*e* = Σ_*i*_ *o*_*i*_*/g*_*0*_), and Pearson’s *χ*^*2*^ statistic can be calculated by the usual approach: *χ*^*2*^ = Σ(o - *e)*^*2*^*/e*. Under ideal conditions, Pearson’s *χ*^*2*^ would approximate the analytical *χ*^*2*^ distribution with its mean and standard deviation equal to the degrees of freedom (*df*) and 2*df*, respectively. While this could be used to make an analytical hypothesis test (as above), in practice there will often be large deviations between Pearson’s *χ*^*2*^ and the analytical distribution, due to violation of the assumptions when expected counts are low (See Supplementary Materials, Figure S2). Instead, we simulate a null distribution of 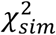 values under the null hypothesis by using permutation within each lineage and recalculating 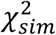 for each replicate. The *p*-value is then equal to the proportion of the 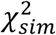 values that exceed the observed *χ*^*2*^ (using all lineages simultaneously), while the effect size is calculated as the mean *C*-score across all pairwise contrasts (simulating 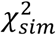 for each pairwise contrast):

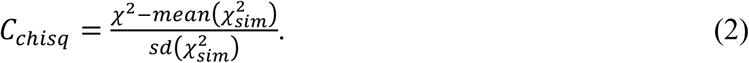

The magnitude of *C*_*chisq*_ therefore represents deviation between the observed amount of repeatability and that expected under the null hypothesis, which will vary as a function of the diversity constraints affecting the trait evolution, but not the number of lineages being compared. While *C*_*chisq*_ relies upon simulation of a null distribution, it can be calculated relatively quickly. Importantly, the magnitude of *C*_*chisq*_ varies linearly with *C*_*hyper*_ (Figure 3A & B), showing that it represents the extent of diversity constraints in the same way as the analytically precise *C*_*hyper*_. While this approach provides a more accurate *p*-value for comparisons of multiple lineages, there is no particular reason to use *C*_*chisq*_ rather than *C*_*hyper*_ for binary input data, as both effect sizes are calculated on a pairwise basis. The main reason that we develop this approach is to extend it to continuously distributed data, which can allow greater sensitivity and avoid arbitrary choices necessary to categorize the commonly used metrics of local adaptation (*e.g*. *F*_ST_ or *p*-values) into “adapted” or “non-adapted”.

**Figure 3.**
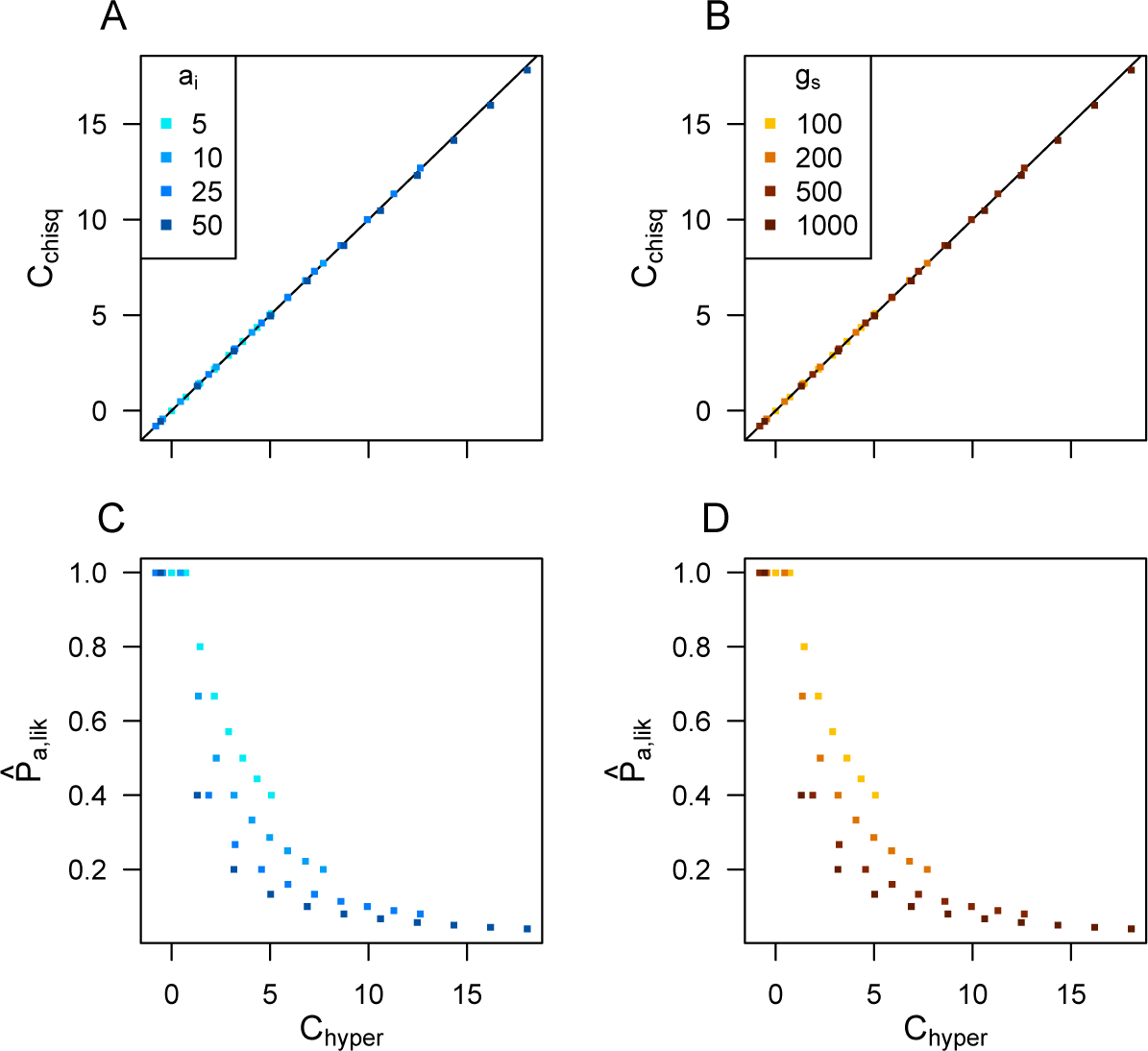
*C*_*chisq*_ and *C*_*hyper*_ provide approximately equal estimates of the magnitude of the diversity constraints driving repeatability, while 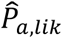 provides an estimate of the proportion of all genes that could potentially contribute to adaptation, which is not collinear with the *C*-scores. Plots show values calculated for simulated datasets generated by randomly drawing two arrays with *g*_*s*_ genes, with *a*_*i*_ loci adapted in one array and *a*_*i*_ + 20 in the other, and then sorting a proportion of the rows in each array to artificially generate more repeatability than would occur by chance (with a different proportion sorted in each replicate). In Panel A&C, *g*_*s*_ = 200; in panel B&D, *a*_*i*_ = 10; 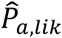 calculated using equation 4.

### Quantifying diversity constraints with continuous data

In many empirical contexts, genome scans for selection yield continuously distributed scores representing the strength of evidence for each locus contributing to adaptation (e.g., *F*_ST_, *p*-values, Bayes factors). Using the same notation as above, but with *α*_*i,j*_ representing the continuous score for the *i*^th^ gene in the *j*^th^ lineage, the total score for each gene can be calculated as a sum across lineages, 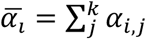, while the mean score over all genes and lineages is 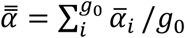. A statistic analogous to the above *χ*^*2*^ can then be calculated as 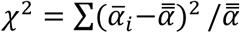, and the same approach for calculating the null distribution of this statistic can then be used to calculate *C*_*chisq*_ according to equation 2. With continuous data, there are additional complexities that arise depending on the distribution of the particular input metric being used and how its magnitude represents evidence for a gene’s involvement adaptation. One approach, which we used in all examples here, is to transform data so that values scale positively and approximately linearly with the weight of evidence for adaptation, by standardizing data within each lineage by subtracting their observed minimum and dividing by their observed maximum, such that the values within each lineage are bounded from 0 to 1. This reduces differences among lineages in the absolute magnitude of metrics representing adaptation, which can be desirable when they vary across many orders of magnitude (e.g. *p*-values of 10^−^10^^ and 10^−^20^^ both provide strong evidence of adaptation). However, if some lineages actually have stronger signatures of adaptation at more loci, then this kind of standardization should not be used, as it would obscure these true differences among lineages. In this case, it would be preferable to use the same standardization across all lineages by subtracting the minimum and dividing by the maximum values observed across all lineages. While Pearson’s *χ*^*2*^ statistic was designed for discrete data, the above approach using continuous data represents the variability among lineages in the same way, as a variance among genes in the sum of their scores representing putative adaptation. The *C*_*chisq*_ statistic on continuous data behaves similarly to the *C*_*hyper*_ statistic across wide ranges of parameter space, as both are formulated in terms of deviations from the null distribution (see below).

### What proportion of the genome can potentially contribute to adaptation?

While the number of genes that potentially contribute to adaptation (*ga*_*s*_) can be estimated using the hypergeometric equation, 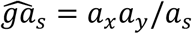, it is difficult to apply this to comparisons of multiple lineages, as some pairwise contrasts may have no overlap in the genes contributing to adaptation (*a*_*s*_ = 0), making the equation undefined. To estimate 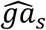 from all lineages simultaneously, we can instead formulate a likelihood-based approach where the probability that we observe locus *i* adapted in *o*_*i*_ lineages is:

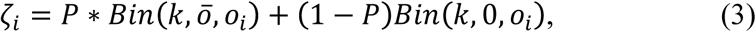

where *Bin*(*n,y,x*) is the probability under the binomial distribution of getting *x* successes in *n* trials, each with probability *y*. As above, *o*_*i*_ is the number of adapted genes in *k* lineages (with *o*_*i*_ Σ_*j*_ *α*_,*i, j*_), *P*_*a*_ is the proportion of *g*_0_ that can actually contribute to adaptation (*P*_*a*_ = *ga*_*s*_ / *n*_*s*_), and *ō* is the probability of each gene contributing to adaptation *ō* = Σ *o*_*i*_/(*ga*_*s*_ *k*). The estimated value of 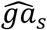 is then the value at which the likelihood function:

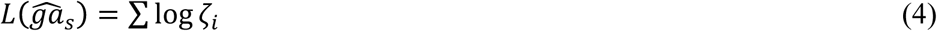

is maximized. Once the maximum-likelihood value of 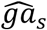 is estimated, this can be expressed either as an absolute number representing the effective number of genes that can contribute to adaptation or as a proportion of the total number of shared genes in the genome: 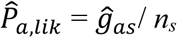. This approach implicitly assumes that all genes that have the potential to contribute to adaptation (*ga*_*s*_) have approximately equal probabilities of actually contributing to adaptation. In very extreme cases, such where one gene is very highly repeatable while other genes only contribute to adaptation in a single lineage, 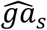 will tend to represent the contribution of the repeatable genes and discount the contribution of the idiosyncratic genes (see Supplementary Materials). Multi-class models could be developed to estimate *ga*_*s*_ for different classes of genes in such scenarios by accounting for their different probabilities of contributing to adaptation (See supplementary materials for a link to scripts containing functions for the above calculations).

## Results

### Comparison between metrics for quantifying convergence

The *C*_*hyper*_, *C*_*chisq*_, and 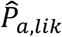 estimators capture different aspects of the biology underlying convergence than other previously used estimators of repeatability. To estimate the repeatability of evolution, Conte et al. [1] used the additive and multiplicative Proportional Similarity (PS_*add*_ and PS_*mult*_) metrics of [30] in a meta-analysis of QTL and candidate gene studies, while Bailey et al. [2] used the Jaccard Index to quantify patterns in bacterial evolution experiments. The PS metrics are defined as 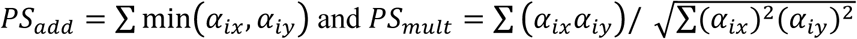, where α_*ix*_ and α_*iy*_ are the relative contribution of gene *i* to adaptation in lineages *x* and *y* [1], while the Jaccard index is defined as *J*= (*A*_*x*_ ∩ *A*_*y*_)/*(A*_*x*_ ∪ *A*_*y*_), where *A*_*x*_ and *A*_*y*_ are the sets of adapted genes in each lineage [2]. Both of these metrics are based on standardizing the number of overlapping adapted loci by the total number of adapted loci, and neither includes information about non-adapted genes that potentially could have contributed to adaptation.

To illustrate the differences between these various metrics of convergence, we generated four example datasets showing either randomly drawn complements of genes with adapted mutations (Figure 4A) or highly convergent datasets drawn from a smaller (Figure 4B) or larger (Figure 4C & D) pool of genes that potentially contribute to trait variation (*g*_*s*_), with differing numbers of loci contributing to adaptation. Scenario C is the most constrained, as it exhibits the same amount of overlap as B, but this overlap is drawn from a larger pool of mutations so it is less likely to occur by chance. While neither the Jaccard index nor the PS metrics distinguish between the B, C, and D scenarios (as the same proportions of genes are being used for adaptation, so repeatability is the same), both the *C*_*chisq*_ and *C*_*hyper*_ metrics show the highest scores for scenario C, because it has the smallest probability of occurring by chance if all genes had equal probabilities of contributing to adaptation. The 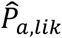 metric also identifies scenario C as most constrained in terms of the smallest proportion genes potentially contributing to adaptation. The 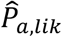 metric also shows that this proportion is equal for scenarios B & D, despite differences in the probability of the observed repeatabilities occurring by chance (as per the *C*-scores). More generally, while 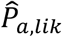 tends to decrease with increasing *C*- score, these metrics differ in magnitude (Figure 3C & D), as they represent different aspects of diversity constraints. In summary, the Jaccard and PS metrics quantify the proportion of genes used for adaptation that are used repeatedly, the C-score metrics represent the probability of the observed repeatability occurring if there were no constraints, and 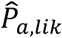 represents the proportion of genes in the genome that are available for adaptation, given the existing diversity constraints (also see Figure S3 for further comparisons).

**Figure 4.**
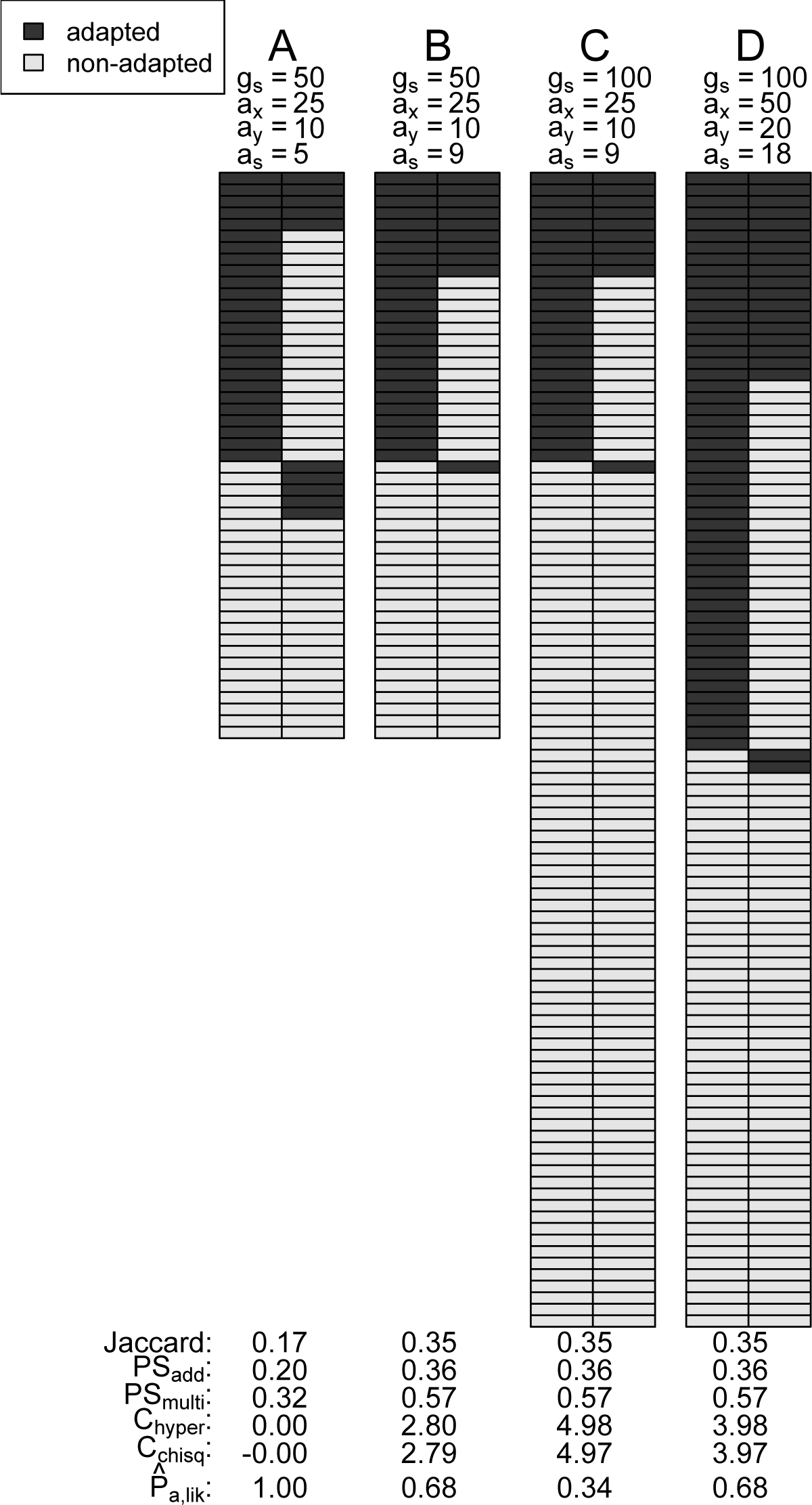
Four example datasets showing different levels of convergent adaptation and a comparison of different metrics assessing overlap among adapted genes. Scenario A is unconstrained and exactly equal to the mean expectation under a random draw; scenarios B & C show the same amount of overlap (*a*_*s*_) and number of adaptively mutated genes (*a*_*i*_), but scenario C is drawn from a larger number of potential genes (*g*_*s*_). Scenario D has the same proportion of overlap as B & C, but twice as many adapted genes.

### Simulating convergence using individual-based simulations

To further explore the effect of population genetic parameters on the behaviour of the above metrics of repeatability and constraint, we used Nemo (v2.3.45; [31]) to simulate two scenarios of two-patches under migration-selection balance: (i) constant size of mutational target with variable proportions of small- and large-effect loci; and (ii) constant number of large-effect loci and variable number of small effect loci, resulting in a variable size of mutational target. For scenario (i), simulations had *n* = *g*_*s*_ = 100 loci, of which *u* loci had alleles of size +/- 0.1, while (100 – *u*) loci had alleles of size +/- 0.01 (with subsequent mutations causing the allele sign to flip from positive to negative or the reverse). For scenario (ii), simulations had 10 large-effect loci with alleles of size +/- 0.1 and *v* small-effect loci with alleles of size +/- 0.01, resulting in a variable size of mutation target. In all simulations, migration rate was set to 0.005 and the strength of quadratic phenotypic selection was 0.5, so that an individual perfectly adapted to one patch would suffer a fitness cost of 0.5 in the other patch (patch optima were +/- 1; similar to [32]). Simulations were run for 50,000 generations and censused every 100 generations. For binary categorization of the input data, loci were considered to be “adapted” if *F*_ST_ > 0.1 for >80% of the last 25 census points (these cut-offs are somewhat arbitrary, but qualitative patterns were comparable under different cut-offs); for continuous input data, raw *F*_ST_ values were used. Results are averaged across 20 runs, each with 20 replicates, with *C*_*chisq*_ calculated across the 20 replicates within each run.

These scenarios further illustrate the difference between the Jaccard and PS_*add*_ metrics of repeatability and the *C*-score and 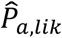 metrics of constraint. In both scenarios, the small effect loci do not tend to contribute much to adaptation because large effect loci are more strongly favoured under migration-selection balance [33], which results in low GF-redundancy. In scenario (i), all metrics show qualitatively similar patterns, with decreasing repeatability occurring as a result of the decreasing constraints that occur as the number of large-effect loci increases, increasing the GP- and GF-redundancy (Figure 5A). By contrast, in scenario (ii), the Jaccard and PS_*add*_ metrics indicate that roughly the same amount of repeatability is occurring regardless of the number of small effect loci and total size of mutational target (Figure 5B). However, over this same range of parameter space, the *C*-score metrics show that constraint increases as the total mutational target is increasing. This occurs because while a larger number of potential routes to an adaptive phenotype are available with increasing number of small effect loci, only the same small number of loci are actually being involved in adaptation (*i.e*. the large effect loci), which is illustrated by the decrease in the 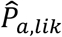 metric. While there are many potential genetic routes to adaptation that could involve these small effect loci (high GP-redundancy), the large effect loci tend to be favoured and repeatedly involved in adaptation (low GF-redundancy). Thus, when the size of the mutational target increases in scenario (ii), the repeatability tends to stay about the same (Jaccard and PS_*add*_) but the amount of constraint is higher (*C*-scores), because a smaller proportion of the available routes to adaptation are being used (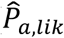). The continuous and binary *C*_*chisq*_ metrics are broadly similar across these parameters because there is very little variation in *F*_ST_ among loci within the same size class (see Supplementary Materials for additional simulations under varying allele effect sizes).

**Figure 5.**
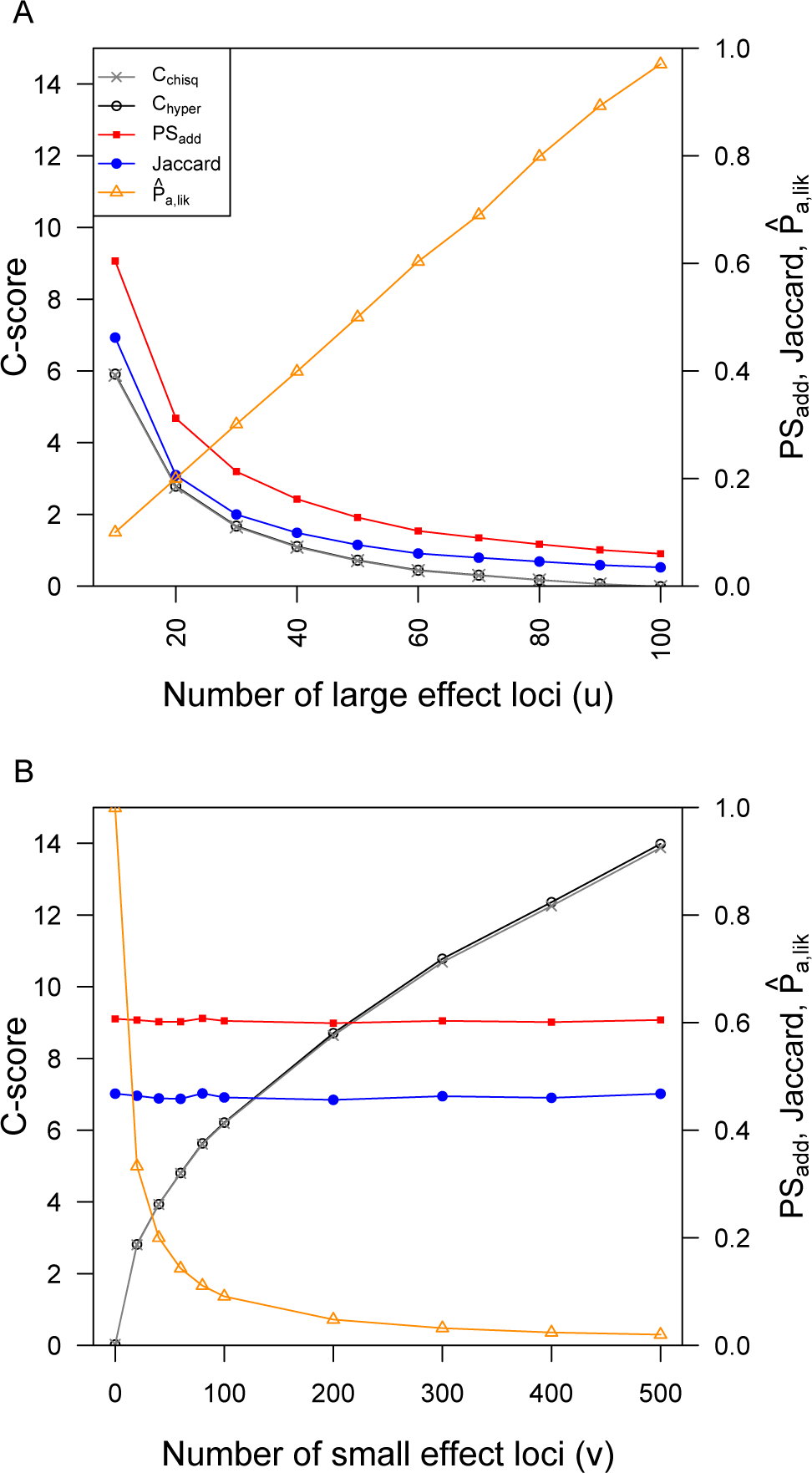
*C*-score metrics of constraint are qualitatively similar to Jaccard and PS_*add*_ metrics of repeatability when simulations have a constant size of mutational target (A), but differ when simulations vary in the size of mutational target (B). 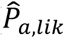 shows qualitatively similar patterns to the *C*-scores, with a decreasing proportion of the genome accessible to adaptation occurring in scenarios with higher *C*-scores and higher constraints. In panel A, all runs have *n*_*s*_ = *g*_*s*_ = 100 loci, with *u* large effect loci and (100 – *u*) small-effect loci. In panel B, there are 10 large-effect loci, and *v* small-effect loci. In both scenarios, simulations were run with *N* = 10,000 individuals in each patch, recombination rate of *r* = 0.5 between loci, and per-locus mutation rate = 10^−^5^^. The calculation of *C*_*hyper*_ is based on categorizing genes as adapted when *F*_ST_ > 0.1, while the calculation of *C*_*chisq*_ is based on *F*_ST_ standardized by subtracting the minimum value and dividing by the maximum within each lineage.

### Adjusting for incomplete sampling of the genome

The amount of constraint quantified by the *C*-score will depend upon the proportion of the mutational target (*g*_*s*_) that is sampled by the sequencing approach, which should be proportional to the sampling of the total number of genes in the genome (*n*_*s*_). Some approaches, such as targeted sequence capture, will sample on only a subset of the total number of genes in the genome, which will therefore cause a bias in the estimation of constraint due to this incomplete sampling, even if the genes included are a random subset of *g*_*s*_. This can be most clearly seen in the calculation of *C*_*hyper*_, where multiplying all the variables in Eq. 1 by a given factor will cause a change in the magnitude of the effect size. By contrast, the Jaccard and PS measures of repeatability are not affected by incomplete sampling. If binary input data are being used and the proportion of *g*_*s*_ that has been sampled can be accurately estimated (*q*), then the calculation of *C*_*hyper*_ can be corrected by dividing all input variables by *q* prior to calculation, yielding a corrected score *C*_*hyper-adj*_. If continuously distributed input data are being used, then the dataset can be adjusted by adding *g*_0_ (1 - *q*) new entries to the dataset by randomly sampling genes with replacement from the existing dataset, and then calculating applying Eq. 2 to this extended set.

To explore the effect of incomplete sampling of the genome on the calculation of *C*-scores and the impact of these types of correction, we constructed a test dataset by concatenating 5 replicates from the simulations in Figure 5A with 10 large effect loci, yielding a dataset with 500 loci in total and a high amount of repeatability. We then sampled a proportion *q* of this total dataset to simulate incomplete representation of the genome and used the above approach calculate uncorrected and corrected *C*-scores. While incomplete sampling can cause considerable bias in *C*-scores, as long as *q* is not too small, these approaches yield relatively accurate corrections of these estimates (Figure 6). At very low values of *q*, the variance in estimation among replicate subsets increases as a result of sampling effects when only a small number of adapted loci are included, but on average the magnitude of the corrected *C’*-score is independent of *q*.

**Figure 6.**
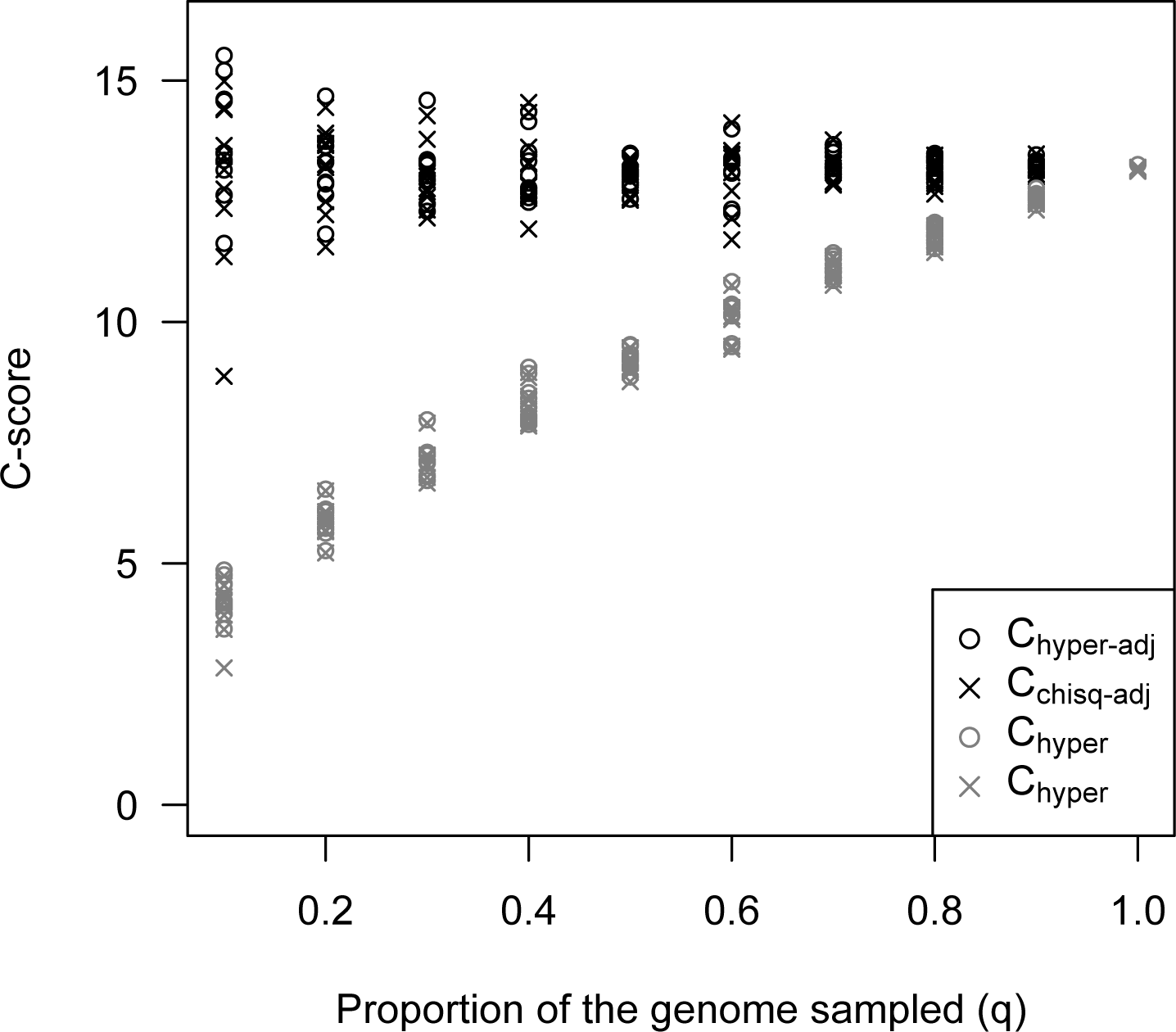
Incomplete sampling of the genome causes a bias in the estimation of *C*-scores (*C*_*hyper*_ and *C*_*chisq*_), but this can be adjusted by using a correction factor (*C*_*hyper-adj*_) or resampling from the existing dataset up to the estimated genome size (*C*_*chisq-adj*_). These approaches yield unbiased *C*-scores, although the variance of the estimates increases due to sampling effects when the proportion of sampled genes (*q*) is small. Figure shows estimates for 10 replicate subsamples performed for each value of *q*.

### Example: Antibiotic resistance in yeast

Experimental evolution studies provide a controlled framework to test theories on the genetic basis of adaptation under a diversity of scenarios. Gerstein et al. (2012) previously conducted an experiment to examine the diversity of first-step adaptive mutations that arose in different lines initiated with the same genotypes in response to the antifungal drug nystatin [34] and in response to copper [35]. The design allowed them to directly test how many different first-step solutions were accessible to evolution when the same genetic background adapted to this same environmental stressor. In the nystatin- evolved lines they identified 20 unique and independently evolved mutations in only four different genes that act in the nystatin biosynthesis pathway: 11 unique mutations in *ERG3*, 7 unique mutations in *ERG6*, and 1 unique mutation in each of *ERG5* and *ERG7* [34]. The genotypic basis of copper adaptation was broader, and there were both genomic (SNPs, small indels) and karyotypic (aneuploidy) mutations identified. If we consider just the genomic mutations, mutations were found in 28 different genes, with multiple mutations identified in four genes (12 unique mutations in *VTC4*, four unique mutations in *PMA1*, and 3 unique mutations in *MAM3* and *VTC1*). If we assume that all genes in the genome could potentially contribute to adaptation (*i.e. g*_0_ = 6604), then *C*_*hyper-nystatin*_ = 32.5, while *C*_*hyper-copper*_ = 12.3, and p < 0.00001 in both cases.

If we assume much lower GP-redundancy and that only the observed genes could possibly contribute to the phenotype (*i.e. g*_0-nystatin_ = 4, *g*_0-copper_ = 28), we can test whether the mutations are still more clustered than expected within these sets. Using the methods outlined above, we find *C*_*hyper-nystatin*_ = 0.35, *p* = 0.002, and *C*_*hyper-copper*_ = 0.43, *p* < 0.0001, indicating that even under the severe developmental-genetic constraints to diversity represented by this model, these data are slightly more overlapping more than expected at random, likely due low GF-redundancy and potentially gene-specific differences in mutation rate. (Because these experiments were initiated using isogenic strains, standing variation was precluded).

Experimental evolution studies lend themselves nicely to future hypothesis testing about the impact of constraint on the genetic basis of adaptation, and provide us with hypotheses about differences between the genes that were and were not observed in the screen. For example, we parsed the Saccharomyces Genome Database (http://www.yeastgenome.org) for genes that have been annotated as “resistance to nystatin: increased”, where this phenotype is conferred by the null mutation. This should be a conservative dataset, as we also expect there could be mutations in additional genes that do not result in a loss-of-function phenotype that could also confer tolerance to nystatin (although we expect that the mutations we recovered in ERG3, ERG5 and ERG6 are all similar to loss of function mutations, ERG7 is inviable when null [34]). This identified an additional five genes (*KES1, OSH2, SLK19, VHR2, YEH2*). We can test whether the five genes without an observed mutation have a negative pleiotropic effect when null, or are in areas of the genome with a lower mutation rate compared to the ERG genes (particularly compared to *ERG3* and *ERG6*). Similar experiments could also be conducted with different *Saccharomyces cerevisiae* genetic backgrounds, with closely related species, or under slightly different environmental conditions (e.g., increased or decreased levels of stressor) to directly examine how different aspects of the genomic and ecological environments influence the observed level of constraints acting on adaptation

### Example: Cold tolerance in conifers

Lodgepole pine and interior spruce both inhabit large ranges of western North America and display extensive local adaptation, with large differences in cold tolerance between northern and southern populations in each species. Recent work studied the strength of correlations between population allele frequencies and a number of environmental variables and phenotypes in each species [36]. Taking one representative environmental variable as an example, a total of 50 and 121 single-copy orthologs showed strong signatures of association to Mean Coldest Month Temperature (MCMT) in pine and spruce, respectively, with 5 of these genes overlapping (based on binary categorization using the binomial cutoff “top candidate” method, as per [36]). This study included a total of 9891 one-to-one orthologs with sufficient data in both species (*i.e.* at least 5 SNPs per gene), so observing 5 genes overlapping corresponds to *C*_*hyper*_ = 5.6 and *p* = 0.00034 under the null hypothesis that all genes had equal potential to contribute to adaptation. Alternatively, it is also possible to estimate *C*_*chisq*_ on continuously distributed data by calculating top candidate scores for each gene using the binomial probability of seeing *u* outliers when there are *v* SNPs in a given gene, with an overall rate of *w* outliers per SNP (as per [36], this yields an index rather than an exact probability, due to linkage among SNPs). This approach is more sensitive to weak signatures of adaptation that occur below the binary categorization cutoff, yielding *C*_*chisq*_ = 5.1 and *p* < 0.00001. Assuming that the 9891 studied genes represent a random sample from approximately 23,000 genes in the whole genome and ignoring divergence in gene content between species (*n*_*s*_ = *n*_*x*_ = *n*_*y*_), the adjusted *C*-scores are *C*_*hyper-adj*_ = 8.6 and *C*_*chisq-adj*_ = 7.8 (with resampling of 50 replicates and 10,000 permutations per replicate), providing a very rough estimate of the total diversity constraints driving repeatability. These diversity constraints correspond to an effective adaptive target of *ga*_*s*_ = 1462 genes that could potentially contribute to adaptation in these species (out of 9891), which yields 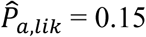. However, a large number of the 50 and 121 genes identified using their “top candidate test” were likely false positives, because there were no controls for population structure during the association test, as this was subsequently accounted for by the among-species comparison. Thus, if we assume a 50% false positive rate for *a*_*x*_ and *a*_*y*_, then *ga*_*s*_ declines to 370 genes, with 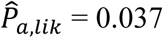. In their analysis, Yeaman et al. [36] used another more sensitive test (null-W) to identify loci with signatures of convergence that were not detected based on overlap in the top candidates lists, which suggests the true amount of repeatability may be higher than *C*_*hyper*_ = 5.6. This example illustrates how these kinds of statistics may be used to make inferences about constraints, but also highlights the sensitivity of the results to small changes in parameters.

## Discussion

*Mc1r* provides perhaps the most well known case of convergent local adaptation at the gene scale, and has been implicated in driving colour pattern variation in mice, lizards, mammoths, fish, and a range of other organisms ([21,37–40]). Extensive study in mice has revealed that over 50 genes can be mutated to give rise to variation in colour pattern [41], yet *Mc1r* consistently tends be one of the main contributors to local adaptation in colour pattern in many vertebrates. Given the apparent high GP-redundancy, this can be seen as a likely example of GF-redundancy constraining the diversity of forms: *Mc1r* has minimal pleiotropic side effects [39,41] and it can mutate to similar phenotypes through numerous different changes in its protein sequence [21,41] and therefore may have a higher rate of mutation to beneficial alleles than other genes with similar per-nucleotide mutation rates. If all of these features of *Mc1r* serve to facilitate adaptation by repeated change in the same gene, should this really be called “constraint”? Again, it is critical to differentiate between “constraints to adaptation” and “constraints on the diversity of forms”. Evolution via *Mc1r* can clearly occur quite readily, and is therefore relatively unconstrained in terms of the potential for contributing to fitness gains over time. But the lack of other equally suitable alternative routes for realizing change in colouration phenotypes constitutes a clear constraint on the diversity of forms. Other genes are more constrained than *Mc1r*, resulting in evolution overall being constrained to evolve similar forms through similar genetic routes. This of course brings up the fascinating question: why do such constraints exist? Are the factors that constrain adaptation actually themselves evolving adaptively?

Our aim here has been to develop a method to quantify the constraints that drive repeatability so that we can test hypotheses about the nature of these constraints on the diversity of forms and reasons they exist. We have focused on how convergence can be inferred from observational studies of local adaptation and experimental investigations of short-term laboratory evolution, but phylogenetic studies of convergence in nucleotide substitution rates could also be used (*e.g*. [42,43]). With any type of study, comparative approaches examining the same trait across different branches of the phylogeny may allow us to infer rates of evolution in constraints and study whether adaptation drives such changes. Comparisons across traits within lineages will illuminate how different kinds of traits (e.g. morphological, physiological, behavioural) are constrained, and whether low GF- and GP-redundancy constitute important constraints at different levels of biological organization (e.g. pathways and products, tissues, organs, integrated traits). Similarly, it will be interesting to examine whether the types of constraint that predominate depend upon critical population genetic parameters such as effective population size (N_e_) that affect the long term efficiency of selection on the developmental/genetic/ecological landscape that gives rise to constraint. While we have focused on repeatability at the gene level, this framework could be applied at other levels of organization, such as gene network, protein domain, or individual nucleotide (reviewed by [3]), and could include the contribution of intergenic regulatory regions if it is possible to identify orthology.

### Hypothesis testing to identify the factors that constrain diversity

Under the simplest null hypothesis that there are no diversity constraints, *g*_0_ = *g*_*s*_ = *ga*_*s*_ = *n*_*s*_ (*i.e.* all genes can give rise to variation in the trait). While simplistic, this approach provides an intuitive method to assess whether the amount of convergence observed is more than expected due to pure randomness. But what do we learn if we reject such a simple null hypothesis? Two inferences can be drawn in this case: many of the genes flagged by our tests for selection are likely evolving by natural selection (*i.e*. they are not all false positives) and some kind of constraint is involved in shaping this adaptation. The former inference means that analyzing comparative data for convergence can provide a powerful tool for identifying the genes involved in local adaptation, as this is often as significant methodological hurdle in evolutionary biology (*e.g*. [36]). The latter inference may seem a straw-man, as few molecular biologists would advocate a model where every gene can mutate to give rise to variation in a given trait. However, different forms of the “universal pleiotropy” model have been assumed in theoretical quantitative genetics [44], and the recently proposed “omnigenic model” advocates extensive pleiotropy [45]. Regardless of whether this null hypothesis represents a plausible biological scenario, it provides a benchmark against which we can quantify how all factors constraining the diversity of forms combine to drive repeatability, which is useful for interpreting patterns of repeatability among species and traits.

In order to make inferences about the potential importance of different kinds of diversity constraints driving repeatability, it is necessary to specify more realistic models for the evolution of local adaptation that incorporate different assumptions about size of the mutational target of the trait, extent of shared standing variation, differences in mutation rate among genes, distribution of mutation effect sizes, and species demography. The simplest modification to the above null model is to represent the extent of GP-redundancy by specifying the number of loci that potentially contribute to trait variation as a subset of the total number of loci in the genome (*g*_0_ *= g*_*s*_ < *n*_*s*_). In the context of eq. (1), reducing *g*_0_ increases both the mean and standard deviation of the hypergeometric distribution and therefore decreases *C*_*hyper*_ and the inferred level of residual (unexplained) constraints. If empirical estimates of *g*_*s*_ result in *C*_*hyper*_ ∼ 0, then it is reasonable to conclude that low GP-redundancy is mainly responsible for the observed amount of convergent adaptation. This would not discount the importance of natural selection overall, as selection on the phenotype is still responsible for adaptation, but would suggest that individual loci are more or less interchangeable and GP-redundancy is high. However, as we have few (if any) conclusive estimates of *g*_*s*_ in highly polygenic traits [46,47], the extent of constraint arising through low GP-redundancy will be difficult to assess without further directed study. Although they are by no means simple experiments to conduct, it should be possible to estimate *g*_*s*_ from QTLs identified in multiple mutation accumulation experiments, as the number of loci detected across all experiments should asymptote towards *g*_*s*_, and rarefaction designs could be used to estimate *g*_*s*_ based on the overlap between QTLs detected in two experiments (although this would likely still be biased by failing to detect loci of small effect). A similar approach could be taken using Genome Wide Association Studies on standing variation for a given trait and comparing the loci identified in different species to assess the proportion of shared loci. Depending on the difference between the size of the mutational target (*g*_*s*_) and the inferred size of the effective adaptive target (*ga*_*s*_), it is possible to draw conclusions about the relative importance of GP- vs. GF-redundancy (Table 1).

**Table 1:**
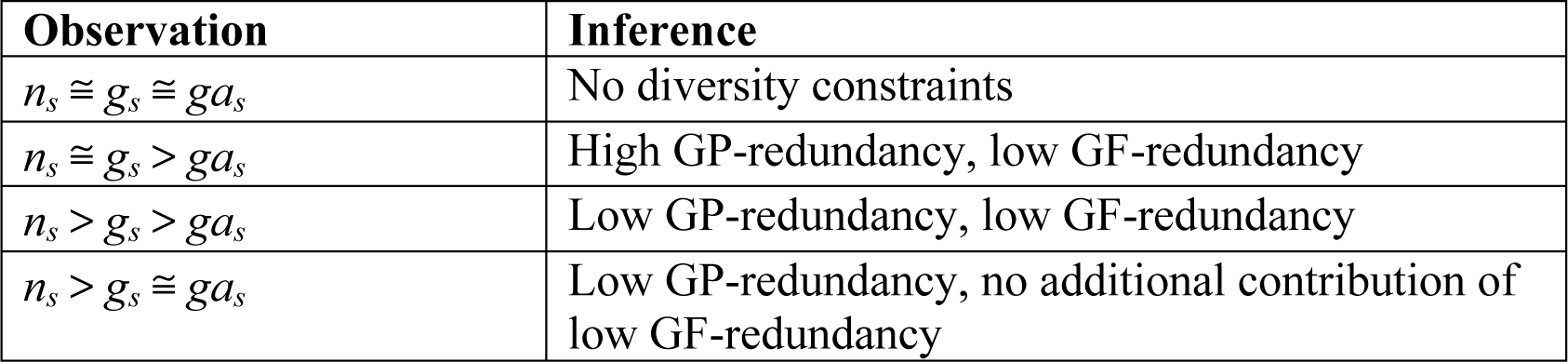
Drawing inferences about the nature of constraints to diversity that drive repeatability

In order to draw inferences about the importance of these types of redundancy, it is critical to account for other factors unrelated to GP- and GF-redundancy that might drive repeatability, mainly through differences among genes in mutation rate or standing variation. The simplest approach to control these factors is to design studies that preclude shared standing variation, either through experiments founded from isogenic strains (*e.g.* [2,34]) or comparisons of distantly related lineages (divergence time >> 4*N*_*e*_) where lineage sorting has been completed (as per [36]). While repeatability could still be driven by differences among genes in mutation rate, this can be seen as a component of redundancy and therefore as factor that can also constrain diversity. By contrast, the existence of shared standing variation occurs mainly due to historical contingency, and is therefore a bias affecting estimation of C-scores, rather than a constraint. As such, parsing the contribution of mutation rate to the repeatability assessed by C-scores and 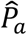 is less critical than parsing the contribution of standing variation when using these as overall metrics of constraint. Unfortunately, in studies of recently diverged natural populations, it is not possible to preclude shared standing variation, so C-scores and 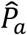 could be strongly driven by this factor and therefore not particularly representative of diversity constraints. The recently developed likelihood-based method for discriminating between convergence via *de novo* mutation, migration, or shared standing variation ([17]) may provide a means to parse these contributions to repeatability and refine the inference of constraint. While testing the null hypothesis of no constraints is relatively straightforward, discriminating among other potential factors constraining diversity is much more complicated. Although it is possible to make very intricate models with variable mutation rates or shared standing variation, selection coefficients or indices of pleiotropy, and other factors determining the likelihood of each gene contributing to adaptation [3,20,26,48], it may be very difficult to actually confidently discriminate between such models.

### Practical considerations in implementation

The accuracy of these metrics will depend critically on the correct identification of the genes contributing to adaptation. Studies of local adaptation are particularly prone to false positives when population structure is oriented on the same axis as adaptive divergence, and it is unclear how extensively methods that correct for population structure induce false negatives or fail to accurately control false positives [49,50]. Assuming false positives are distributed randomly throughout the genome in each lineage, failure to remove them will cause the *C*-scores derived here to be biased downwards. Failure to identify true positives (*i.e.* false negatives) will impair the accuracy of *C*_*chisq*_ but will not necessarily bias it in one direction or the other, as this would depend upon the underlying biology. Assuming false negatives are randomly distributed in the genome, they could reduce the magnitude of *C*-scores due to lower information content. But on the other hand, because large-effect loci are more likely to be both detected and convergent and small effect loci are more likely to be missed, false negatives will tend to bias *C*-scores upwards. As it is typically necessary to set arbitrary cutoffs for statistical significance to identify putatively adapted loci, we might expect *C*_*chisq*_ to increase with increasing stringency of these cutoffs, as this would be expected to reduce false positives. However, as there are many potential contingencies and interactions between the factors that affect these two types of error, there is a clear need for both theoretical studies on how the repeatability of local adaptation is affected by the interplay between demography and selection (*e.g*. [26]), and refinement of these methods to derive confidence intervals taking into account likely error rates.

A particularly important problem to address in implementing this method is that false positives may be non-randomly distributed throughout the genome in a similar way in different lineages. As local variations in the rate of mutation or recombination can drive genome-wide patterns in some metrics used to identify selection and adaptation [51–53], this could lead to signatures of convergence among distantly-related species if such patterns are conserved over long periods of evolutionary time. For example, genome-wide patterns of variation in nucleotide diversity, *F*_ST_, and *d*_xy_ were all significantly correlated across three distantly related bird species, likely driven in part by conservation of local recombination rate coupled with linked selection [54]. The extent of convergence of local recombination rates appears to vary considerably among species [55–58], so it will be important to consider this factor as a potential driver of similarity in the genomic signatures used to identify selection. Methods for identifying signatures adaptation that are explicitly linked to a phenotype or environment of interest across multiple pairs of populations may be less likely to be affected by such factors, as recombination and linked selection are unlikely to drive a pattern of repeated correlation between allele frequency and phenotype/environment. However, such methods are still vulnerable to potential biases that arise from the complex interplay between genomic landscape, selection, and recombination, and further study in both theoretical and empirical contexts will be important to test the robustness of different methods to this important source of bias.

While studying adaptation across multiple pairs of populations can greatly increase the power to detect signatures of selection when all populations are adapting via the same loci, such methods are inherently unable to detect idiosyncratic patterns where different populations of a given species are adapting via different loci. By its very nature, it may be very difficult, if not impossible to detect local adaptation in traits with high GP- or GF-redundancy, as each pair of populations may be differentiated via a different set of loci [32]. If local adaptation is much more readily detected when it arises repeatedly within a lineage, then it will be difficult to identify conclusive cases with low C-scores, causing an overestimation of the prevalence of highly repeated adaptation.

If patterns of genomic convergence are compared among multiple differentially-related lineages, it is important to consider their phylogeny when testing the importance of phylogenetic sharing of different factors affecting the propensity for gene reuse [59]. Also, the ability to resolve orthology relationships decreases with increasing phylogenetic distance, which can affect the estimation of *n*_*s*_. Similarly, the set of genes in a trait’s mutational target (*g*_*i*_) is expected to evolve over time, so the set of shared genes should decrease with phylogenetic distance (so that *g*_*i*_ – *g*_*s*_ increases with divergence time), leading to decreased repeatability over time [1]. When studies include multiple differentially-related lineages, it is probably useful to estimate *C*-scores on both a pairwise and mean-across-all-lineages basis to more clearly describe cases where convergence is high within pairs of closely related lineages but low among more distantly related lineages.

Finally, physical linkage is a factor that could critically affect the measurement of repeatability, as neutral alleles in other genes linked to a causal allele will tend to respond to indirect selection, causing spurious signatures of selection/local adaptation. If the same causal gene is driving adaptation in two lineages, this will tend to overestimate repeatability on a gene-by-gene basis, whereas the opposite will occur if different causal genes are driving adaptation. Yeaman et al. [36] found significantly elevated levels of linkage disequilibrium (LD) among candidate genes for local adaptation, which may have arisen due to physical linkage (with or without selection on multiple causal loci) or statistical associations driven by selection among physically unlinked loci. In this case, the fragmented genome and lack of suitable genetic map precluded a comprehensive analysis of the impact of LD. If genome/genetic map resources permit, it may be possible to analyse repeatability on haplotype blocks rather than individual genes, which could minimize the biases due to physical linkage.

### Comparison to other metrics of repeatability

A large number of metrics have been developed to characterize similarity among ecological communities, which can be broadly grouped based on binary vs. quantitative input data and whether they account for joint absence of a given type (reviewed in [60]). In most cases, these metrics are not derived from a probability-based representation of expectations, though Raup and Crick [61] quantified an index of similarity based on the *p*-value of a hypergeometric test (see also [62]). The *C*_*hyper*_ metric that we have developed here uses the same underlying logic as the Raup-Crick metric, but quantifies the effect size as a deviation from the expectation under the null hypothesis in units of the standard deviation of the null distribution. The *C*-score and 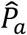 metrics developed here provide a complement to metrics of repeatability that have been used in previous studies of convergence at the genome scale (*e.g*. [1,2]). Whereas the Jaccard, PS_*add*_, and other similar metrics represent how commonly a given gene tends to be used in adaptation, the *C*-score metrics quantify how much constraint is involved in driving this observed repeatability, whereas 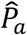 quantifies the proportion of the genome that is effectively available for adaptation. In some cases, these metrics will be qualitatively similar in quantifying patterns of convergence (e.g. Figure 5A), but in other cases they will diverge considerably, because the *C*-scores are explicitly aimed at representing the importance of genes that could contribute to adaptation but do not.

## Conclusions

When adaptation uses the same genes, does this occur because there are only a few genetic ways to make a given phenotype, or because only a few ways are optimally fit? Direct quantification of genotype-phenotype landscapes that could answer this question has been conducted in some very specific cases, yielding fascinating insights into how such landscapes shape evolution (*e.g*., [63]). Unfortunately, for most traits in most species, it is currently impractical to systematically and directly quantify the mapping of genotype to phenotype to fitness. However, studying the repeatability of adaptation can provide an indirect way to learn about the topology of this landscape. In this case, a first step towards quantifying how the genotype-phenotype-fitness map shapes adaptation is to compare the genes that potentially contribute to standing variation (*g*) with the genes that potentially contribute to adaptation (*ga*). The recently formulated “omnigenic model” posits that most genes can have some effect on phenotypic variation in a given trait but that they can be subdivided into “core” vs. “peripheral” genes with large and small effects, respectively [45]. Other reviews of Genome-Wide Association Studies have come to similar conclusions about the large size of mutational target and interchangebility of allelic effects underlying standing variation [64,65]. These studies therefore imply that many complex traits exhibit high GP-redundancy. But what fraction of the alleles contributing to standing variation actually have the potential to contribute to adaptation? Is GF-redundancy also high? Given that most phenotype-affecting mutations are thought to be deleterious rather than beneficial [66], it seems likely that most of the alleles contributing to standing variation are simply deleterious variants that have not yet been purged from the population, and therefore may not be representative of the “stuff” of long-term adaptation. This is probably especially true for the genetic basis of disease in humans and other animals, which in most cases are deleterious by definition. Studying the repeatability of adaptation at the genome scale will provide insights into the extent of GF-redundancy by comparing the change in metrics of repeatability for the null hypothesis of no constraints (complete GP-redundancy) vs. a null hypothesis under a more limited mutational target (lower GP-redundancy). As we now have many examples of convergent adaptation at different levels of organization [1,3,67], it seems likely that many traits exhibit quite limited GF-redundancy and that standing variation does not necessarily correspond to long-term adaptive potential. We hope that the methods formulated here provide a useful way to compare results among studies and test these hypotheses directly. Unfortunately, it will be much easier to detect the cases with high convergence than to conclusively demonstrate non-convergence, so a great deal of care will be required when interpreting these data.

## Acknowledgments

We would like to thank J. Mee, D. Schluter, and the Yeaman lab for comments and discussion during the preparation of this manuscript. Special thanks to L. Harris for suggesting Pearson’s *χ*^*2*^ when S.Y. & A.G. were mired in combinatorics. This work was supported by AIHS and NSERC Discovery grants to S.Y., a Genome Canada grant to S.Y. and M.C.W., a CIHR Banting Postdoctoral Fellowship to A.C.G., an ARC grant to K.H., and an NSERC Discovery Grant to M.C.W. The project was enabled in part by computational support provided by WestGrid and Compute Canada.

